# Dysregulated miR-3959-3p in response to Lumpy skin disease virus

**DOI:** 10.1101/2023.02.13.528427

**Authors:** Sakshi Pandita, Davinder Singh, Naveen Kumar, Yogesh Chander

## Abstract

Lumpy skin disease virus (LSDV), a member of the *Capripoxvirus* genus, causes substantial economic losses in the livestock industry and is rapidly spreading among various LSDV-free countries across the globe. Viral infections are known to alter the cellular miRNA expression profile of the host significantly.Besides being important biomarker candidates, circulating miRNAs have a significant role in controlling viral infection and antiviral immune responses, including several molecular mechanisms. miR-3959-3p, a significantly downregulated miRNA revealed in RNA-sequencing results of LSDV-infected LT cells, was selected to bedetected in the serum of LSDV-infected and uninfected cattle sera (40 LSDV-infected and 40 LSDV-uninfected). We optimized quantitative real-time PCR (qRT-PCR) for quantitative determination of miR-3959-3p in the bovine sera samples. The level of miR-3959-3p appears to be towards lower side in the LSDV-infected as compared to the uninfected animals. However, no significant correlation could be established between the two groups due to fluctuations in the miRNA levels in both groups. This is the first report on the detection of circulating miRNA in LSDV-infected cattle sera.The dysregulation pattern of miR-3959-3p appears to suggest that several other miRNAs need to be explored and may serve as biomarkers for LSDV infection. However, this needs further investigation by screening several other miRNAs and on large number of LSDV positive and negative animals.

**Author summary:** MicroRNAs are the key regulators of viral infections. However, in LSDVinfection the miRNA response is greatly unknown.

In this study, miRNA expression in Vero cell linepost LSDV infection was studied for the first time.

One of the miRNAs identified in the RNA-sequencing results i.emiR-3959-3p, was shown to be downregulated LSDV infection. We detected the levels of mir-3959-3p in sera of LSDV-infected and uninfected cattle to explore its potential as a biomarker.

## Introduction

Lumpy skin disease virus (LSDV) is one of the three viruses in the genus *Capripox*virus within the Poxvirus family and is responsible for Lumpy skin disease, one of the most economically significant disease of domestic ruminants in Africa and Asia [1]. LSD is also known as “Pseudo-urticaria”, “Neethling virus disease”, “knopvelsiekte” and “exanthema nodularis bovis” [2]. This highly infectious, generalized and often fatal disease results in huge production losses and chronic debility among livestock. [3]. Owing to its rapid spread and devastating economic impacts, World Organization for Animal Health (OIE) has listed LSD as a notifiable disease LSDV mainly affects cattle, however, mild disease has been observed in buffalo population. Due to the detection of viral DNA in skin nodules and anti-LSDV antibodies in convalescent blood samples, confirmation of moderate LSDV infection in camels, horses, and deer has been observed (unpublished). This implies that LSDV has the potential to infect other host species, prompting the inclusion of these species in the epidemiology and management of LSDV infection in animals.

The main clinical feature of LSDV includes slightly raised and firm skin nodules of 2-7 cm in diameter [4]. Other clinical signs include fever, enlarged lymph nodes, nasal discharge, and salivation, keratitis, pneumonia [5]. Sometimes edema is seen in the limbs which subsequently results in lameness [6]. The reproductive complications of the disease include, abortion, orchitis and mastitis [7]. The disease is more severe in lactating cows and there is a sharp drop in milk production [8]. The evidence of in-contact susceptible animals showing neutralizing antibodies is suggestive of the fact that subclinical infection may be present in some animals during an LSDV outbreak [9]

Transboundary animal diseases (TADs) like LSD are a serious threat to sustainability and socio-economic status of livestock [10]. In 1929, Zambia reported the first LSDV outbreak. [11]. Subsequently, between the years 1943 and 1945, more LSDV outbreaks were reported in Botswana, Zimbabwe and the Republic of South Africa till it further spread to the African continent [2]. In the Middle East, LSDV outbreak was reported in Oman around the year 1984 [12]. Later, Egypt, Israel, Bahrain, Yemen, United Arab Emirates also reported the outbreak of LSDV [12]. Re-emergence of outbreaks is not uncommon as many countries have LSDV outbreaks after the first reported outbreaks [13]. The Asian countries like India, Bangladesh, China, Nepal, Vietnam, Myanmar, Sri Lanka, Thailand, Malaysia have recently reported LSDV outbreaks which raises a matter of serious concern for the livestock and dairy sector [14]. India witnessed its first outbreak in the second week of August in 2019 in the state of Orissa which was followed by two more outbreaks in the same month which affected 79 cattle. An overall morbidity and mortality of 8.48% and 0% respectively were observed in the country [15]. However in the year 2022, India witnessed a drastic increase in LSD cases especially in the western state of Rajasthan and Gujarat with high mortality [16]. According to some preliminary observations, the high pathogenicity of LSDV/2022 strain which is circulating in Southeast Asia is likely due to its ability to cause severe nodule formation in visceral organs, in particular lungs [16]. The main source of LSDV outbreaks worldwide are illegal movement of animals and the vectors which transmit the virus [17].

MicroRNAs (miRNAs) are short non-coding RNAs (ncRNAs) of 22 nucleotides that play important roles in regulating gene expression[18].The two main roles of miRNA include gene regulation and intercellular signalling, both of which play a significant role in viral infections[19].miRNAs in whole blood and plasma are potential for enhancing disease diagnosticsand recent influx of data suggests that circulating miRNAs in serum are considered as promising candidates as diagnostic biomarkers in veterinary medicine[20]. Therefore, the major aim of this study is to analyse the miRNA response to LSDV-infected and uninfected cattle sera.

**Fig. 1:**
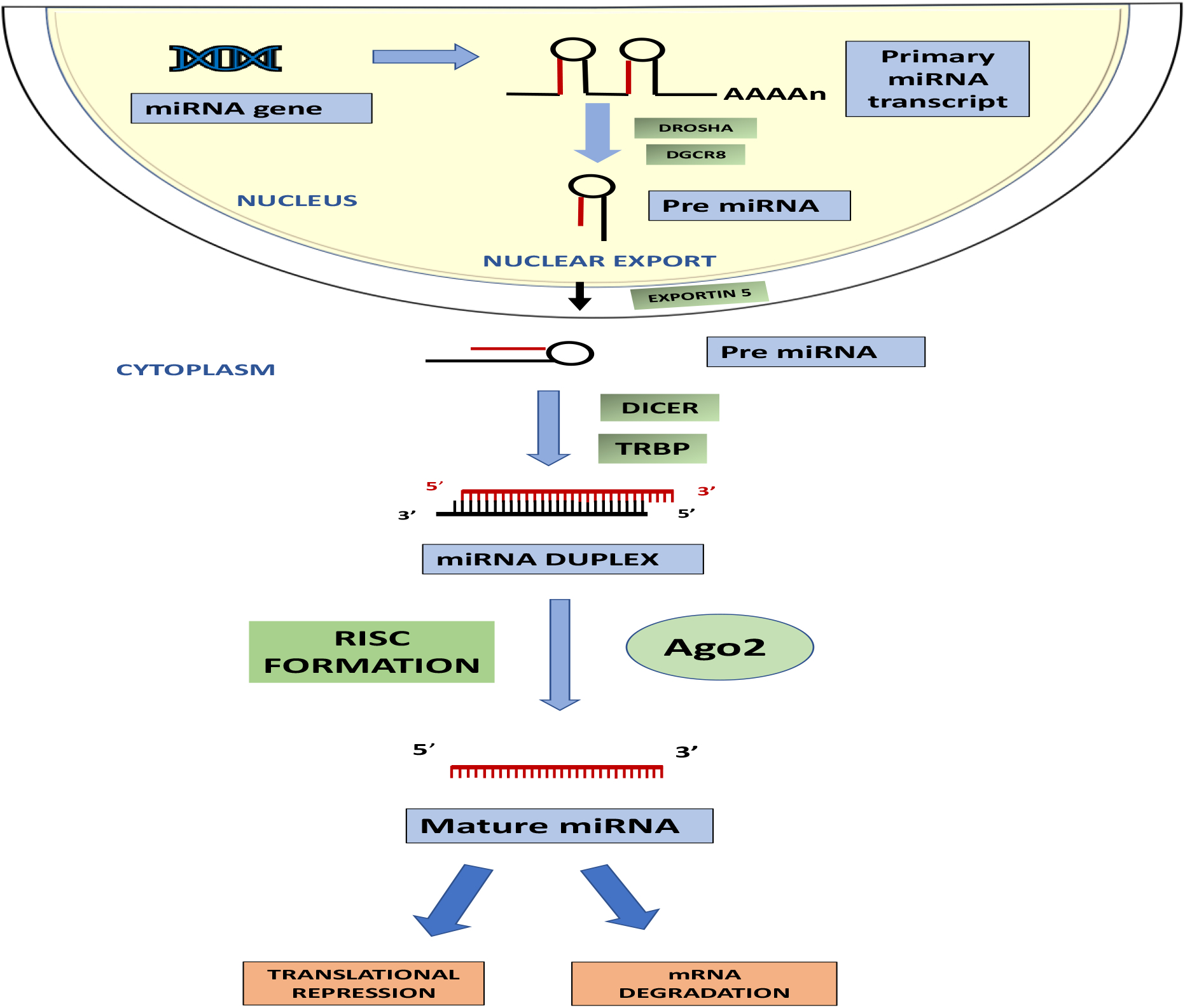
miRNA biogenesis: miRNAs are processed from the primary transcript using a two-step sequential mechanism involving two RNase III nucleases. The primary precursor (pri-miRNA) is processed into an approximately 70 nucleotide long stem-loop structure by nuclear RNase III Drosha present in the microprocessor complex. The resultant pre-miRNA is exported to the cytoplasm by a complex of Exportin-5. The final maturation of miRNA occurs with the help of Dicer, another RNase III nuclease that processes the pre-miRNA into a 22 bp double-stranded RNA. The processing is often coupled with the formation of the ribonucleoprotein complex known as RISC (miRNA-Induced Silencing Complex). The RISC minimally consists of one strand of the miRNA (called “guide strand”) in addition to Dicer, TRBP, PACT and Argonaute (Ago) proteins.

## Results

### miRNA response in naturally infected animals

Circulating miRNAs present in animal serum have been explored in various viral diseases. Their high stability in serum makes them potentially interesting candidates for diagnostic and other clinical uses. Out of the several miRNAs identified to be significantly dysregulated in miRNA profiling studies, we measured the levels of one of the highly dysregulated (downregulated) miRNA (miR-3959-3p) in sera of LSDV infected or uninfected (naïve) cattle. The miR-3959-3p was detectable in sera of both LSDV-infected and uninfected animals. Although, the levels of miR-3959-3p appeared to be on the lower side in the LSDV-infected cattle, however, due to high individual variations, no significant difference could be observed in the expression level of miR-3959-3p in the sera of LSDV-infected and uninfected cattle.

**Fig 2:**
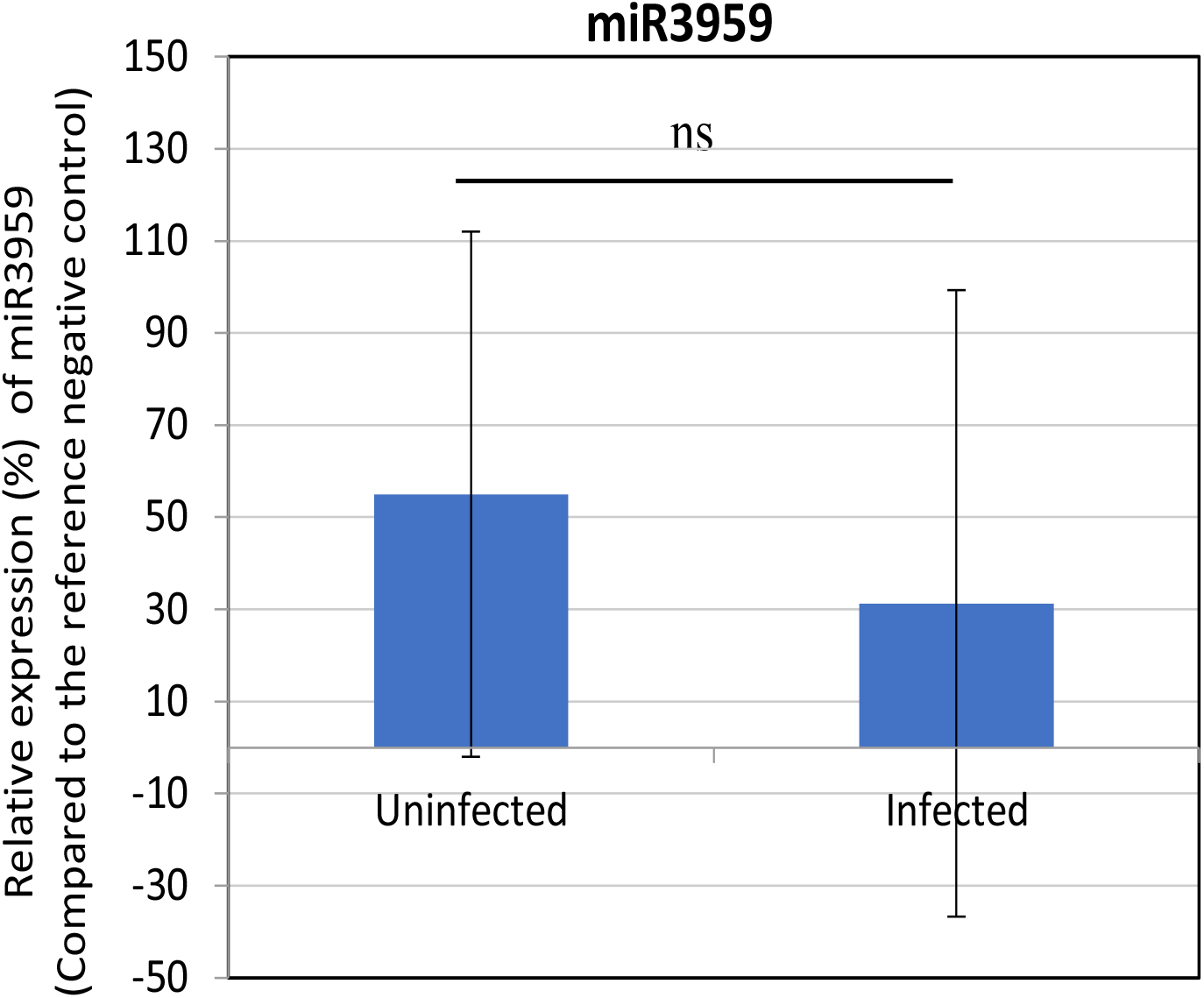
miR-3959-3p level in the sera of LSDV infected and uninfected animals. Serum samples (n=40) were collected from the cattle that had a recent history of LSDV infection. Equal number of serum samples (n=40) were also collected from cattle that had no history of LSDV infection and were free from antibodies against LSDV. 200 µl of the serum sample from each animal was subjected to RNA isolation, its quantitation and normalization to have an equal amount of final RNA concentration in each sample. This was followed by poly (A) tailing, cDNA synthesis, pre-amplification of cDNA and quantitative determination of miR-3959-3p levels by qRT-PCR. Five naïve animals that were never exposed to LSDV were kept under controlled conditions were used as reference uninfected control. The relative differences in the level of miR-3959-3p miRNA between LSDV-infected and uninfected animals were analysed by using student’s t-test. The error bars indicate standard deviation. ns=Non-significant difference.

## Discussion

LSDV is a serious transboundary disease of livestock and its recent emergence in Asia and the Indian subcontinent has made this viral disease a serious concern among the livestock owners. The economic consequences of LSDV can be disastrous. For instance, it is estimated that with 10.5 percent of the cattle and buffaloes at risk in Asia, direct losses of USD 1459 million could be caused by this deadly virus [21].

India reported its first LSDV outbreak in the state of Odisha in the year 2019 and gradually, in a span of just a few years, witnessed a devastating LSDV outbreak in 2022 in 15 states affecting over 2 million animals and accounting for 100,00 deaths [15, 16]. The fact that LSDV has recently emerged in Asia, combined with the evidence that such outbreaks in the past likely result in the disease being endemic in the affected area, highlights the importance of extensively studying the deadly virus and its epidemiology [22]. A viral disease spreading among the cattle population of the world’s highest milk-producing country at such an alarming rate makes timely diagnosis and prevention an immediate need. Virus isolation, electron microscopy and Polymerase chain reaction (PCR) are the common diagnostic techniques used to identify the virus. A widely accepted rapid and sensitive method of detecting LSDV includes real-time PCR [23] All *Capripoxviruses* cross-react serologically, thus serology lacks the ability to distinguish between animals that have received vaccinations with the LSDV, SPPV, or GTPV strain vaccines from those that have not [24].The virus can last a long time under ambient circumstances, for example, it can survive in necrotic nodules for 33 days, crusts of dehydrated skin for 35 days, and air-dried hides for at least 18 days [24]. Therefore, skin biopsies are collected for diagnosis of the disease. However, collecting skin biopsies on a large scale during an outbreak can be a tedious process. On the other hand, studies have evidenced that blood samples are less reliable for detecting the viral genome due to the short span of viremia, which shows positive results in a short window of about 4-11 days post-infection [25]. Studies have also demonstrated that some cattle with LSDV infection or vaccination may not show seroconversion [26]. Additionally, detection of the virus during the incubation period and in sub-clinical animals is most likely not possible. Such cases play a significant role in the transmission of the virus, and there is a substantial gap in the study of such claims. Virus neutralisation test has been validated but is slow, expensive, and has a low sensitivity [27]. The biology and pathophysiology of LSDV are poorly understood. Therefore, exploring the host-LSDV virus relationship will significantly increase our understanding of the disease progression and lead to novel, easy, and quick diagnostic methods.

Serum is a vital biomarker detection candidate as compared to biopsy samples, especially in veterinary science, where handling animals is a tedious task. Circulating miRNAs in serum have been detected in various viral diseases in animals [28-30]. In this study, no significant difference in the levels of miR-3959-3p could be observed between the sera derived from LSDV-infected and uninfected cattle, partly because of large individual animal variation. The miRNAs can be dysregulated in response of various abiotic and the biotic factors [31]. One potential explanation is that the animals in the field would have been exposed to other concurrent infections [32], and therefore, necessitates conducting such studies under controlled conditions. Since all the animals were not of identical genetic makeup (not pure line), they would have responded differently [33]. Furthermore, since all the sera samples were not derived from a single geographical location, the different abiotic/management/nutritional/environmental factors would also have also affected the miRNA levels in infected sera [31, 33, 34]. In addition, since the time window (time after infection) of collection of all the LSDV positive serum was not identical, this would have also affected the temporal expression of miRNA, as has been observed by previous workers [28-30] Nevertheless, a previous study on Hendra virus (HeV) infection also suggests that miRNA detection in field samples is complicated, partly because of missing out the time window for detecting a particular miRNA [35]. Therefore, it can be concluded that circulating miRNAs in viral disease as biomarkers are a bit challenging and this gap can further be explored by conducting a well-designed case-control studies in diverse animal populations.

## Materials and methods

### Cells

African green monkey kidney (Vero cells) and primary lamb testicular cells (LT cells) were available at NCVTC, Hisar.

### Virus

Vero cell adapted lumpy skin disease virus (LSDV/2019/India/Ranchi/P50, GenBank Accession OK422494), originally isolated from an LSD affected cattle from an outbreak in Ranchi (2019) was available at NCVTC, ICAR-NRCE, Hisar [9]. The virus was quantified by determination of tissue culture infective dose 50 (using TCID_50_) as described previously [9]

### Propagation and quantitation of LSDV

The virus was grown and titrated in Vero cells by determination of the tissue culture infective dose 50 (TCID_50_) as per the standard procedure described previously (Kumar et al. 2021). For bulk culture, Vero cells were infected at MOI 0f 0.1. The virus was harvested at 72 hpi, a time point when >70% cells showed the CPE. The viral titres were ∼10^7^ TCID_50_/ml in most instances.

### Cell culture medium and reagents

Dulbecco’s modified *Eagle’s* Medium (DMEM) supplemented with 10-15% foetal bovine serum (FBS) (Sigma, St. Louis, USA) and antibiotic solution (Sigma, St. Louis, USA) was used to culture primary lamb testicle- and Vero cells. The confluent monolayers of cells were trypsinized by 0.25% Trypsin-EDTA solution (Sigma Aldrich, St. Louis, USA).

Primary lamb testicular cells (LT cells) were infected with LSDV virus at multiplicity of Infection (MOI) of 5 for 2 hours (h), followed by washing with PBS and addition of fresh medium.The mock-infected and virus-infected cells were collected at 12, 48 and 72 hpi. Total RNA was isolated from Mock-infected and LSD infected cells by TRIzol™ (Invitrogen, USA) and RNA was quantified byNanoDrop spectrophotometry(Thermo Fisher, Wilmington,DE, USA). Two micrograms of total RNA isolated from Mock-infected, and LSDV infected lamb testicle (LT) cells be sent for quantitative and qualitative determination of all the possible miRNAs (miRNA profiling) at Next Generation Sequencing (NGS; Illumina) platform. miRNA deep sequencing was outsourced at Wipro Pvt Ltd. Bengaluru (India).

NGS was analysed for identification of the dysregulated (upregulated/downregulated) miRNAs and miR-3959-3p selected for detection in the serum. Serum samples from LSDV-positive animals (n=40) and LSDV-negative animals (n=40) were available at NCVTC. Serum samples from animals that tested positive in PCR for both the LSDV genome in skin nodules and for antibodies against LSDV in serum neutralization test (SNT) were included in the study as positive samples. Inversely, the samples collected from asymptomatic animals that tested negative in SNT were considered as negative samples.

The desalted primers were synthesized by Sigma-Aldrich chemicals Pvt. Ltd., Bangalore. As directed by the manufacturer, these primers were diluted to make stock solution (1:10) using nuclease-free water. To create the working primer solution, the stock solution was further diluted (1:10). The primer pairs used in the PCR reactions are as follows: Forward-5’ TGTATGTCAACTGATCCACAGTCC 3’ and Reverse-5’GGCCACGCGTCGACTAGTAC 3’. 200µl of serum was used for RNA extraction using TRIzol™ (Invitrogen, USA). The RNA was quantified using nanodrop photo spectrometry. This was followed by Poly (A) tailing and cDNA synthesis pre-amplification of cDNA and finally qRT-PCR.

### Poly(A) tailing and cDNA synthesis

Poly (A) tailing of 25 µl of normalized RNA was done using *E. coli* poly (A) polymerase (New England Biolabs^®^ Inc., Ipswich, MA, USA), and the reaction mixture was incubated at 37^0^ C for 30 minutes. This was followed by synthesis of cDNA from the poly(A) tailed RNAs primed by oligo adapter primer (OAP). 20 µl of poly(A) tailed RNA was incubated at 72^0^ C for 10 minutes with 1 µl of oligo adaptor primer (OAP). Total cDNA was synthesized by adding 9 µl of reaction mixture to the OAP primed poly(A) tailed RNA to make a total reaction mixture volume of 30 µl and PCR was carried out under the following thermic profile conditions: annealing at 25^0^ C for 10 minutes followed by extension at 42^0^ C for 50 minutes and finally deactivation at 72^0^ C for 10 minutes.

### Pre-amplification of cDNA

Due to their low concentrations, tiny size, and absence of reference values in biological samples, circulating miRNA analysis is difficult, therefore, reliable miRNA measurement might be made possible by pre-amplifying complementary DNAs [36].

Pre-amplification of the miRNA cDNA enhances the number of detectable miRNAs, boosts the sensitivity of the qPCR array, and preserves the relative levels of miRNA expression. [37]. Therefore, in case of serum samples, where circulating miRNAs had to be detected, pre-amplification of cDNA was carried out using Q5High-Fidelity 2X Master Mix (New England Biolabs^®^ Inc., Ipswich, USA), according to the manufacturer’s instructions. Amplification of the cDNA was done using miRNA-specific forward primers (FP) along with Abridged Anchor Primer (AAP) as the reverse primer (RP). 2 µl of cDNA was used as a template and the thermal cycler conditions included: initial denaturation at 95°C for 10 min, 10 cycles of cyclic denaturation at 95°C for 15 secs, annealing at 60°C for 30 secs, extension at 72°C for 30 secs and final extension at 72°C for 5 mins and final hold at 4°C.

### qRT-PCR

In the present study, quantitative real-time polymerase chain reaction (qRT-PCR) was used to determine the expression of miRNA. The qRT-PCR was performed using SYBR® Green Supermix (Bio-Rad, Hercules, USA). Amplification of the cDNA was done using miRNA-specific forward primers (FP) along with Abridged Anchor Primer (AAP) as the reverse primer (RP). Thermal cycler conditions of initial denaturation at 95°C for 10 mins, 45 cycles of cyclic denaturation and annealing at 95°C for 15 secs and 60°C for 60 secs respectively were used in QuantStudio3 Real-Time PCR System (Thermo Fisher Scientific, Singapore). The miRNA expression levels were calculated using the 2^-ΔΔCt^ method [38]and were normalized to the U6 expression). The qRT-PCR amplification curves were evaluated using QuantStudio Software v1.4.2 (Thermo Fisher Scientific, Massachusetts, USA). The level of significance between the two groups viz., LSDV-infected sera and uninfected sera were analysed using student’s t-test.

## Acknowledgments

A part of this study belongs to MVSc research work of Dr. Sakshi Pandita. Mulatu E, Feyisa A. Review: Lumpy skin disease. J Vet Sci Technol. 2018;9(535):1-8.

